# Whole genome sequence analysis of *Shigella* from Malawi identifies fluoroquinolone resistance

**DOI:** 10.1101/2020.11.20.391235

**Authors:** George E. Stenhouse, Khuzwayo C. Jere, Chikondi Peno, Rebecca J. Bengtsson, End Chinyama, Jonathan Mandolo, Amy K. Cain, Miren Iturriza-Gómara, Naor Bar-Zeev, Nigel A. Cunliffe, Jennifer Cornick, Kate S. Baker

**Affiliations:** University of Liverpool, Liverpool, UK; Malawi-Liverpool-Wellcome Trust Clinical Research Programme, College of Medicine, University of Malawi, Blantyre, Malawi; NIHR Health Protection Research Unit in Gastrointestinal Infections, University of Liverpool, Liverpool, UK; ARC Centre of Excellence in Synthetic Biology, Department of Molecular Sciences, Macquarie University, North Ryde, Australia; International Vaccine Access Center Department of International Health, Johns Hopkins Bloomberg School of Public Health, Baltimore, USA

**Author notes:** These authors contributed equally to this work (Jennifer Cornick, Kate S. Baker).

## Abstract

Increasing antimicrobial resistance and limited alternative treatments led to fluoroquinolone resistant *Shigella* strain inclusion on the WHO global priority pathogens list. In this study we characterised multiple *Shigella* isolates from Malawi with whole genome sequence analysis, identifying the acquirable fluoroquinolone resistance determinant *qnrS1*.

## Introduction

*Shigella* is the second leading cause of diarrhoeal death globally with the greatest disease burdens seen in low and middle-income countries (1). Approximately one third of these deaths are in children under the age of five, with infection potentially causing chronic health effects (1, 2). Antimicrobial resistance (AMR) increasingly limits treatment options, threatening to reverse hard won reductions in diarrhoeal mortality in high-burden areas (3). Meanwhile, vaccines are still in development. There is, therefore, a need for effective public health solutions.

Increasing prevalence of fluoroquinolone resistant (FQR) strains and limited, widely effective alternatives to this first-line treatment, has led to these strains being included on the WHO global priority pathogens list (4). Resistance can be acquired *de novo* through a double mutation in the *gyrA* gene (amino acids 83 and 87) of the quinolone resistance determining region (QRDR), with a third mutation in the *parC* gene (AA80) ameliorating the fitness cost (5). Resistance can also be acquired through horizontal transmission of FQR genes (5).

*Shigella* is reported as a leading cause of diarrhoea among hospitalised children in Malawi, however, there is little information on the circulating strains (6). Whole genome sequence analysis (WGSA) has been successfully applied to investigate *Shigella* epidemiology and AMR determinants, and can greatly aid in disease control in high-burden areas, such as Malawi (7). Here, we have applied WGSA to characterise *Shigella* strains and AMR determinants in Malawi. Providing important baseline information for public health interventions including antibiotic treatment and deployment of *Shigella* vaccines, the development of which is a WHO priority (8).

### The study

All biochemically-confirmed shigellae collected during a rotavirus vaccine evaluation programme between 2012 and 2015, isolated from faecal samples collected from children under the age of 5 hospitalised with acute gastroenteritis at the Queen Elizabeth Central Hospital, Blantyre, Malawi, were subjected to whole genome sequencing (Supplementary) (9). Species were confirmed with a maximum-likelihood phylogeny of the *Shigella*/*Escherichia coli* clade, generated from a core single nucleotide polymorphism (SNP) alignment (40075 SNPs) using quality trimmed reads mapped to a complete reference genome (Supplementary). Only those genomically-confirmed as *Shigella* or *E. coli* (8/10) were included in the study. All *Shigella* isolates were serotyped *in silico* with ShigaTyper (v1.0.6, https://github.com/CFSAN-Biostatistics/shigatyper).

From our eight isolates, four *Shigella* serotypes were identified: two *S. flexneri* 3a (Sf3a, Phylogroup 2), one *S. flexneri* 4av (Sf4av, Phylogroup 7), two *S. flexneri* 6 (Sf6) and one *S. boydii* 2 (Sb, Clade 3). Two isolates were *E. coli* (Figure 1A). The lack of *S. sonnei* isolates in our collection was expected as *S. sonnei*, though highly prevalent globally, is typically associated with high-income nations and industrialisation (10). Prevalence of *S. boydii* in Africa is thought to be low, isolated in 5.7% of cases of a multi-national study (11). However, detection with our small sample size would still be likely by chance (binomial test p-value = 0.375) and may not be due to higher prevalence in Malawi. The strain diversity indicates several, distinct strains circulate in Malawi; further characterisation is needed to aid effective vaccine development for the region.

**Figure 1.**
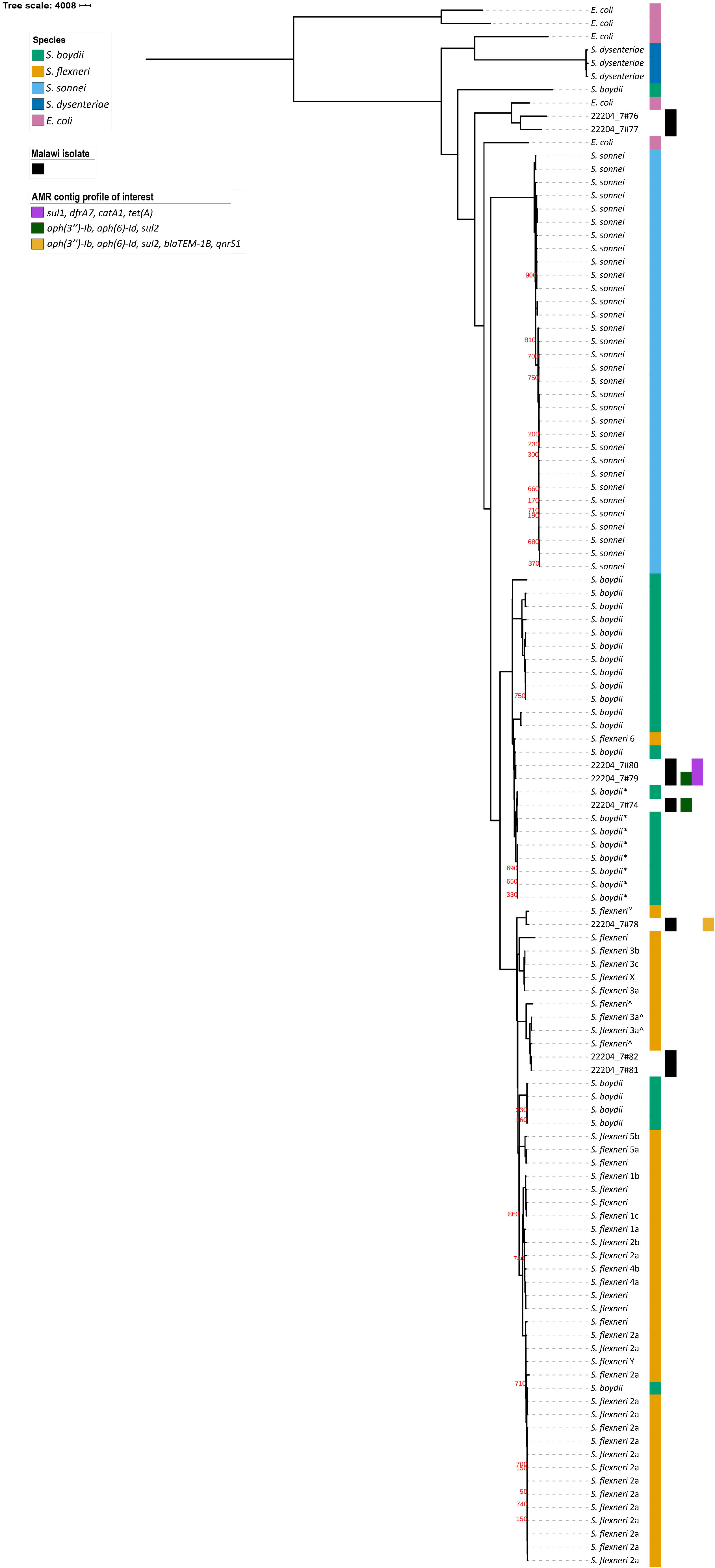
Maximum-likelihood phylogeny of eight isolates from Malawi, contextualised among *Escherichia coli* and *Shigella* and highlighting some AMR profiles of interest. AMR profiles of interest are those of contiguous sequences which are MDR and shared across multiple isolates, and in one isolate carries additional AMR genes including an FQR gene (*qnrS1*). They are indicated by columns to the right of the tree (A) and by coloured bars in AMR profile chart (B). **A.** Columns to the right of phylogeny indicate, from left to right, species, study isolate, AMR profiles of interest. GTR+G substitution model, 1000 bootstrap validation and mid-point rooted. * = *E. coli* clade 3, ^ = *S. flexneri* phylogroup 2, ^γ^ = *S. flexneri* phylogroup 7. All bootstrap values for internal nodes with support <900 are displayed. Scale bar is in SNPs per site. **B.** Intersection of individual AMR genes by isolate and AMR gene profile by contig.

Identification of *E. coli* among the samples is likely due to the close relatedness between the two species. *Shigella* is a specialised pathovar of *E. coli*, sharing a disease phenotype with enteroinvasive *E. coli* (EIEC), that is highly adapted to humans (12). To determine whether the *E. coli* isolates were EIEC we looked for the presence of the mxi-spa locus, found on the large virulence plasmid (pINV). This locus encodes a Type 3 Secretion System and secreted effector proteins which produce the distinctive invasive disease phenotype (12, 13). Contiguous sequences (contigs) from quality-assessed isolate draft-genomes, assembled using Unicycler (v0.4.7) (14), were compared against a reference EIEC pINV to identify locus presence (Supplementary). Using this approach, one *E. coli* isolate was identified as EIEC (99% identity and 100% coverage with mxi-spa locus).

As resistance is a growing hinderance to effective treatment, we determined genotypic AMR profiles for each isolate using starAMR (v0.5.1; https://github.com/phac-nml/staramr; Supplementary). All *Shigella* isolates were predicted to be multi-drug resistant (MDR), carrying genes conferring resistance to three or more drug classes, demonstrating MDR *Shigella* circulate in Malawi (Table 1). While overall resistance was high, we observed limited diversity in AMR genes and predicted resistance profiles; 13 genes encoded resistance to 7 antimicrobial classes (Figure 1B, Table 1). This suggests that there are treatments which remain effective in Malawi, such as azithromycin, though this would need confirming in a larger study and may change with mass drug administration programs.

**Table 1.** Antimicrobial resistance genotypic and predicted phenotypic profiles of Malawian Shigella isolates by contiguous sequence. ^ω^ StarAMR identified IncFIB(K) plasmid, ^Γ^ StarAMR identified MDR IncQ1 plasmid, ^т^ possible multi-copy plasmid.

| Isolate ID | Contig length (bp) | Resistance genes | Predicted resistance drug class |
| --- | --- | --- | --- |
| 22204_7#74 | 9131 <sup>o</sup> | *tet(A)* | Tetracycline |
| (*S. boydii* 2) | 2798 | *aph(3'')-Ib, aph(6)-Id, sul2* | Aminoglycoside, sulfonamide |
|  | 2353 | *dfrA7* | Trimethoprim |
|  | 1831 | *blaTEM-1B* | Aminopenicillin |
| 22204_7#78 | 48082 | *tet(A)* | Tetracycline |
| (*S. flexneri* 4av) | 17334 | *aph(3'')-Ib, aph(6)-Id, sul2, blaTEM-1B, qnrS1* | Aminoglycoside, sulfonamide, aminopenicillin, fluoroquinolone |
|  | 1588 | *dfrA14* | Trimethoprim |
| 22204_7#79 | 34138 | *tet(A), sul1, dfrA7, catA1* | Tetracycline, sulfonamide, trimethoprim, chloramphenicol |
| (*S. flexneri* 6) | 6200 | *aph(3'')-Ib, aph(6)-Id, sul2* | Aminoglycoside, sulfonamide |
| 22204_7#80 | 25733 | *tet(A), sul1, dfrA7, catA1* | Tetracycline, sulfonamide, trimethoprim, chloramphenicol |
| (*S. flexneri* 6) | 6200 | *aph(3'')-Ib, sul2* | Aminoglycoside, sulfonamide |
| 22204_7#81 | 11392 <sup>r</sup> | *dfrA14, sul2, tet(A)* | Trimethoprim, sulfonamide, tetracycline |
| (*S. flexneri* 3a) |  |  |  |
| 22204_7#82 | 45377 | *tet(B)* | Tetracycline |
| (*S. flexneri* 3a) | 8773 | *aadA1, blaOXA-1, catA1* | Aminoglycoside, aminopenicillin, chloramphenicol |
|  | 6790 <sup>r</sup> | *aph(6)-Id, dfrA14, sul2* | Aminoglycoside, trimethoprim, sulfonamide |

Mobile genetic elements (MGE) are important drivers of AMR dissemination, so we explored the genetic context of our AMR genes to look for evidence of them being within MGEs (15). Resistance contigs from two isolates were identified as likely plasmid contigs, also using starAMR (supplementary): one Sf3 isolate carried a MDR IncQ1 plasmid, and the Sf4av isolate carried an AMR IncFIB(K) plasmid (Table 1). Another MDR contig had a read depth 5.29-fold higher than the chromosomal contigs, possible evidence of a multi-copy plasmid (Table 1). Meanwhile, the same AMR gene profile was identified in contigs from multiple isolates across distinct phylogroups (Sf4av and Sf6), possibly indicating an MGE (Figure 1, Table 1). While there was limited evidence, due to the limitations of short read sequencing, taken together, our data supports a role for MGEs in the spread of MDR *Shigella* in Malawi.

To predict FQR, we looked for the presence of either point mutations in the QRDR, or FQR-associated genes. Point mutations were identified by comparing the *gyrA* and *parC* amino acid sequences across all isolates, with amino acid identity at resistance-associated sites confirmed as expected based on the literature. One isolate (Sf4av) was predicted FQR, due to the presence of *qnrS1* (Table 1). A mutation at *parC* R91Q was also identified, however, the same variation was present in all isolates, *E. coli* and *Shigella*, and likely represents natural variation rather than a resistance adaptation. Unfortunately, phenotype data were unavailable to confirm these findings.

Detection of *qnrS1* indicates acquirable FQR is present among *Shigella* in Malawi. As this gene is typically carried on a plasmid, a BLAST comparison of the FQR contig (17334 bases) against the NCBI nt database was performed; more accurate methods of plasmid detection were unavailable due to the short sequence reads (5). Three hits against *S. flexneri* plasmids (KY848295.1, CP024474.1, LR213454.1) were found (e-score = 0.0, query sequence coverage and identity ≥99.9%), suggesting that this gene might be carried on a plasmid. This would mean a high risk of widespread FQR in Malawi and neighbouring regions; further study into the prevalence and nature of FQR in the region is needed.

## Conclusions

Together, the high proportion of MDR *Shigella,* the acquirable FQR and the AMR plasmids detected in this study show that, without intervention, controlling shigellosis in Malawi will be increasingly difficult. Highlighting the importance of further research into the epidemiology of *Shigella* in the region to ensure effective disease control.

## Acknowledgements

GES is supported by a studentship from the MRC Discovery Medicine North (DiMeN) Doctoral Training Partnership (MR/N013840/1). KCJ is supported by a Wellcome International Training Fellowship (201945/Z/16/Z). KSB is supported by a Wellcome Trust Clinical Research Career Development Fellowship (106690/A/14/Z). AKC was supported by an Australian Research Council (ARC) DECRA fellowship (DE180100929). The authors would like to acknowledge the technical assistance of Trevor Wilson and Chikondi Jassi, and the bioinformatic assistance of Caisey Pulford and Lewis Fisher.

Nigel Cunliffe is affiliated to the National Institute for Health Research (NIHR) Health Protection Research Unit in Gastrointestinal Infections at University of Liverpool, in partnership with Public Health England, in collaboration with University of Warwick. The views expressed are those of the author(s) and not necessarily those of the NIHR, the Department of Health and Social Care or Public Health England.

Ethical approval was obtained from the National Health Sciences Research Committee, Lilongwe, Malawi (#867) and the Research Ethics Committee of the University of Liverpool, Liverpool, UK (000490).

---

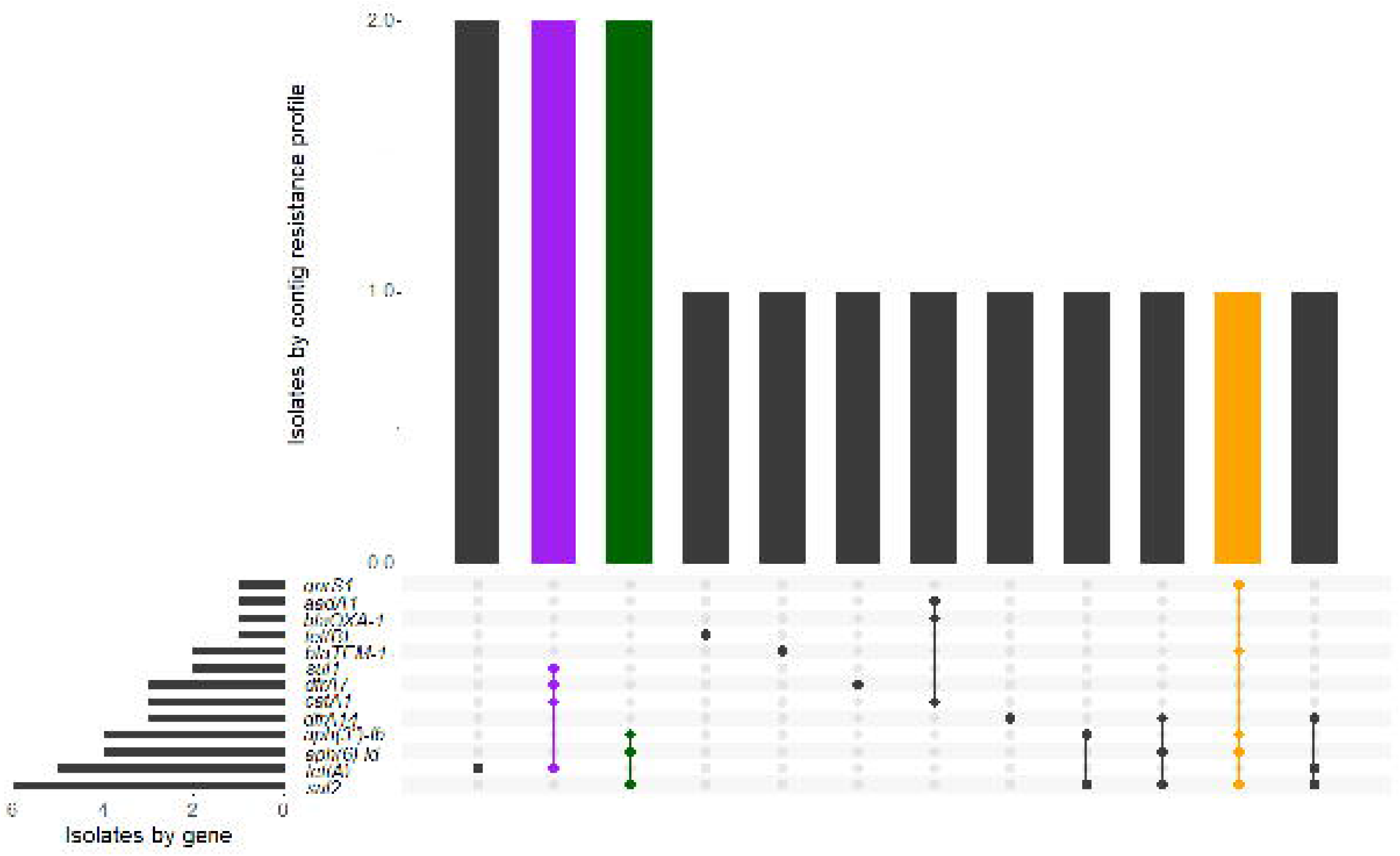

## Notes

### Competing Interest Statement

The authors have declared no competing interest.

