## Supplementary materials for "Whole genome sequence analysis of *Shigella* from Malawi identifies fluoroquinolone resistance"

### Supplementary Appendix

#### Supplementary Appendix Table – Reference and study isolates

| **Accession** | **Database** | **Species and strain** | **Reference** |
| --- | --- | --- | --- |
| CP011417.1 | NCBI Nucleotide | Enteroinvasive *E. coli* plasmid CFSAN029787_01 | (1) |
| NC_004337.2 | NCBI Nucleotide | *S. flexneri* 2a strain 301 chromosome | (2) |
| NC_004851.1 | NCBI Nucleotide | *S. flexneri* 2a strain 301 plasmid CP301 | (2) |
| SRR3237960 | NCBI Short read archive | *S. boydii* | (3) |
| SRR3237961 | NCBI Short read archive | *S. boydii* | (3) |
| SRR3237963 | NCBI Short read archive | *S. boydii* | (3) |
| SRR3237964 | NCBI Short read archive | *S. boydii* | (3) |
| SRR3237965 | NCBI Short read archive | *S. boydii* | (3) |
| SRR3234365 | NCBI Short read archive | *S. boydii* | (3) |
| SRR3234362 | NCBI Short read archive | *S. boydii* | (3) |
| SRR3237801 | NCBI Short read archive | *S. boydii* | (3) |
| SRR3237803 | NCBI Short read archive | *S. boydii* | (3) |
| SRR3237805 | NCBI Short read archive | *S. boydii* | (3) |
| SRR3237806 | NCBI Short read archive | *S. boydii* | (3) |
| SRR3237808 | NCBI Short read archive | *S. boydii* | (3) |
| SRR3237809 | NCBI Short read archive | *S. boydii* | (3) |
| SRR3237810 | NCBI Short read archive | *S. boydii* | (3) |
| SRR3237816 | NCBI Short read archive | *S. boydii* | (3) |
| SRR3237817 | NCBI Short read archive | *S. boydii* | (3) |
| SRR3237837 | NCBI Short read archive | *S. boydii* | (3) |
| SRR3237947 | NCBI Short read archive | *S. boydii* | (3) |
| SRR3237948 | NCBI Short read archive | *S. boydii* | (3) |
| SRR3237949 | NCBI Short read archive | *S. boydii* | (3) |
| SRR3237950 | NCBI Short read archive | *S. boydii* | (3) |
| SRR3237951 | NCBI Short read archive | *S. boydii* | (3) |
| SRR3237953 | NCBI Short read archive | *S. boydii* | (3) |
| SRR3237955 | NCBI Short read archive | *S. boydii* | (3) |
| SRR3237957 | NCBI Short read archive | *S. boydii* | (3) |
| SRR3237959 | NCBI Short read archive | *S. boydii* | (3) |
| SRR3237962 | NCBI Short read archive | *S. boydii* | (3) |
| ERR042803 | NCBI Short read archive | *S. flexneri* 2a | (4) |
| ERR042850 | NCBI Short read archive | *S. flexneri* 2a | (4) |
| ERR048281 | NCBI Short read archive | *S. flexneri* | (5) |
| ERR048288 | NCBI Short read archive | *S. flexneri* | (5) |
| ERR048302 | NCBI Short read archive | *S. flexneri* 2a | (4) |
| ERR048305 | NCBI Short read archive | *S. flexneri* | (5) |
| ERR048317 | NCBI Short read archive | *S. flexneri* | (5) |
| ERR048339 | NCBI Short read archive | *S. flexneri* 2a | (4) |
| ERR126987 | NCBI Short read archive | *S. flexneri* 2a | (4) |
| ERR126993 | NCBI Short read archive | *S. flexneri* | (5) |
| ERR127032 | NCBI Short read archive | *S. flexneri* 1a | (6) |
| ERR127033 | NCBI Short read archive | *S. flexneri* 1b | (6) |
| ERR127034 | NCBI Short read archive | *S. flexneri* 1c | (6) |
| ERR127035 | NCBI Short read archive | *S. flexneri* 2a | (6) |
| ERR127036 | NCBI Short read archive | *S. flexneri* 2b | (6) |
| ERR127037 | NCBI Short read archive | *S. flexneri* 3a | (6) |
| ERR127038 | NCBI Short read archive | *S. flexneri* 3b | (6) |
| ERR127039 | NCBI Short read archive | *S. flexneri* 3c | (6) |
| ERR127040 | NCBI Short read archive | *S. flexneri* 4a | (6) |
| ERR127041 | NCBI Short read archive | *S. flexneri* 4b | (6) |
| ERR127043 | NCBI Short read archive | *S. flexneri* 5a | (6) |
| ERR127044 | NCBI Short read archive | *S. flexneri* 5b | (6) |
| ERR127046 | NCBI Short read archive | *S. flexneri* X | (6) |
| ERR127047 | NCBI Short read archive | *S. flexneri* Y | (6) |
| ERR127048 | NCBI Short read archive | *S. flexneri* strain E1037 | (6) |
| ERR1363976 | NCBI Short read archive | *S. flexneri* 2a | (4) |
| ERR1364007 | NCBI Short read archive | *S. flexneri* 2a | (4) |
| ERR1364014 | NCBI Short read archive | *S. flexneri* 2a | (4) |
| ERR1364050 | NCBI Short read archive | *S. flexneri* 2a | (5) |
| ERR1364087 | NCBI Short read archive | *S. flexneri* 2a | (5) |
| ERR1364097 | NCBI Short read archive | *S. flexneri* 2a | (5) |
| ERR127045 | NCBI Short read archive | *S. flexneri* 6 | (5) |
| ERR1364106 | NCBI Short read archive | *S. flexneri* 2a | (5) |
| ERR1364137 | NCBI Short read archive | *S. flexneri* 2a | (5) |
| ERR200376 | NCBI Short read archive | *S. flexneri* 2a | (5) |
| ERR217085 | NCBI Short read archive | *S. flexneri* | (5) |
| ERR449043 | NCBI Short read archive | *S. flexneri* 3a | (7) |
| ERR449077 | NCBI Short read archive | *S. flexneri* 3a | (7) |
| ERR559526 | NCBI Short read archive | *S. flexneri* 2a | (8) |
| ERR832464 | NCBI Short read archive | *S. flexneri* | (5) |
| ERR832481 | NCBI Short read archive | *S. flexneri* | (5) |
| SRR7886341 | NCBI Short read archive | *S. flexneri* |  |
| ERR024610 | NCBI Short read archive | *S. sonnei* | (9) |
| ERR024611 | NCBI Short read archive | *S. sonnei* | (9) |
| ERR025692 | NCBI Short read archive | *S. sonnei* | (9) |
| ERR025698 | NCBI Short read archive | *S. sonnei* | (9) |
| ERR025701 | NCBI Short read archive | *S. sonnei* | (9) |
| ERR025724 | NCBI Short read archive | *S. sonnei* | (9) |
| ERR025726 | NCBI Short read archive | *S. sonnei* | (9) |
| ERR025737 | NCBI Short read archive | *S. sonnei* | (9) |
| ERR025747 | NCBI Short read archive | *S. sonnei* | (9) |
| ERR025749 | NCBI Short read archive | *S. sonnei* | (9) |
| ERR025751 | NCBI Short read archive | *S. sonnei* | (9) |
| ERR025754 | NCBI Short read archive | *S. sonnei* | (9) |
| ERR025762 | NCBI Short read archive | *S. sonnei* | (9) |
| ERR025765 | NCBI Short read archive | *S. sonnei* | (9) |
| ERR025767 | NCBI Short read archive | *S. sonnei* | (9) |
| ERR025768 | NCBI Short read archive | *S. sonnei* | (9) |
| ERR028673 | NCBI Short read archive | *S. sonnei* | (9) |
| ERR028675 | NCBI Short read archive | *S. sonnei* | (9) |
| ERR028677 | NCBI Short read archive | *S. sonnei* | (9) |
| ERR028679 | NCBI Short read archive | *S. sonnei* | (9) |
| ERR028688 | NCBI Short read archive | *S. sonnei* | (9) |
| ERR028695 | NCBI Short read archive | *S. sonnei* | (9) |
| ERR028700 | NCBI Short read archive | *S. sonnei* | (9) |
| ERR028705 | NCBI Short read archive | *S. sonnei* | (9) |
| ERS127148 | NCBI Short read archive | *S. sonnei* |  |
| ERR200471 | NCBI Short read archive | *S. sonnei* | (10) |
| ERR200544 | NCBI Short read archive | *S. sonnei* | (10) |
| ERR200550 | NCBI Short read archive | *S. sonnei* | (10) |
| ERR212328 | NCBI Short read archive | *S. sonnei* | (10) |
| ERR316322 | NCBI Short read archive | *S. sonnei* | (10) |
| ERR316396 | NCBI Short read archive | *S. sonnei* | (10) |
| ERR319257 | NCBI Short read archive | *S. sonnei* |  |
| ERR1014536 | NCBI Short read archive | *S. dysenteriae* strain M115_cloned | (11) |
| ERR1014032 | NCBI Short read archive | *S. dysenteriae* strain M216 | (11) |
| ERR1013817 | NCBI Short read archive | *S. dysenteriae* strain 98_4962 | (11) |
| ERR1013857 | NCBI Short read archive | *S. dysenteriae* strain IPSP_14_940 | (11) |
| ERR1014544 | NCBI Short read archive | *S. dysenteriae* strain CDC_62_5000_PF1 | (11) |
| ERR1014501 | NCBI Short read archive | *S. dysenteriae* strain CRIE_904 | (11) |
| ERR1014154 | NCBI Short read archive | *S. dysenteriae* strain CAR10 | (11) |
| ERR1014555 | NCBI Short read archive | *S. dysenteriae* strain 13_05279 | (11) |
| ERR1014042 | NCBI Short read archive | *S. dysenteriae* strain SPH_1546 | (11) |
| ERR1014002 | NCBI Short read archive | *S. dysenteriae* strain E44600_86 | (11) |
| SRR10997234 | NCBI Short read archive | *E. coli* strain HS |  |
| SRR1509643 | NCBI Short read archive | *E. coli* strain EDL933 | (12) |
| SRR2061820 | NCBI Short read archive | *E. coli* strain CFT073 | (13) |
| SRR2169510 | NCBI Short read archive | *E. coli* strain ED1a-119 |  |
| SRR2169556 | NCBI Short read archive | *E. coli* strain IAI1-116 |  |
| ERR2525592 | NCBI Short read archive | Malawi Isolate | This study |
| ERR2525594 | NCBI Short read archive | Malawi Isolate | This study |
| ERR2525595 | NCBI Short read archive | Malawi Isolate | This study |
| ERR2525596 | NCBI Short read archive | Malawi Isolate | This study |
| ERR2525597 | NCBI Short read archive | Malawi Isolate | This study |
| ERR2525598 | NCBI Short read archive | Malawi Isolate | This study |
| ERR2525599 | NCBI Short read archive | Malawi Isolate | This study |
| ERR2525600 | NCBI Short read archive | Malawi Isolate | This study |

#### Supplementary Appendix Methods

Whole-genome sequencing was performed at the Wellcome Trust Sanger institute according to in-house protocols (14, 15).

A maximum-likelihood phylogenetic tree was generated from a core-SNP alignment (40075 SNPs) using RAxML-NG (v0.6.6; GTR+G substitution model, 1000 bootstrap validation and mid-point rooted) (16). The SNP alignment was created using snippy (v4.3.6; <https://github.com/tseemann/snippy>) using quality trimmed sequence reads mapped against the complete *S. flexneri* 2a strain 301 genome (chromosome and plasmids, Supplementary Table). All SNPs in plasmids, mobile genetic elements and putatively recombinant regions identified by gubbins (v2.4.1, filter threshold: 31%) were excluded (17).

Quality trimming of sequence reads was performed with Trimmomatic (v0.38) and SeqTK (v1.3, <https://github.com/lh3/seqtk>). Any reads which failed to meet quality standards, visualised with fastQC (v0.11.8), were excluded. This included all the unpaired reverse reads from the Malawi isolates.

A genotypic resistance profile was generated for each isolate by comparing all isolate contigs against the resfinder database using starAMR (v0.5.1; gene identity and overlap ≥99% and ≥90% respectively; <https://github.com/phac-nml/staramr>) (18). All draft genomes were assembled using Unicycler (v0.4.7) and were of sufficient quality, assessed using quast (v8.13) against the *S. flexneri* 301 complete genome (19, 20). Comparison against the plasmidfinder database was used to identify contigs which might be from plasmids (contig identity and overlap ≥98% and ≥80% respectively).

To examine fluoroquinolone resistance, point mutations in QRDR were identified by comparing the *gyrA* and *parC* amino acid sequences across all isolates. Sequences were extracted from the prokka (v1.14.5) annotated genomes and aligned with clustal omega (21, 22). Amino acids at resistance-associated sites were extracted to confirm identity as expected based on the literature.

A BLAST (v2.10.0) search against enteroinvasive *E. coli* pINV (Supplementary Table) was used to identify isolate contigs as possibly encoding the mxi-spa locus. These contigs were then visually compared against the reference pINV, using the Artemis Comparison Tool, to confirm a match against the mxi-spa locus (23).
